# Distinct neural representations of content and ordinal structure in auditory sequence memory

**DOI:** 10.1101/2020.12.05.412791

**Authors:** Ying Fan, Qiming Han, Simeng Guo, Huan Luo

## Abstract

Two forms of information – frequency (content) and ordinal position (structure) – have to be stored when retaining a sequence of auditory tones in working memory (WM). However, the neural representations and coding characteristics of content and structure, particularly during WM maintenance, remain elusive. Here, in two electroencephalography (EEG) studies, by transiently perturbing the ‘activity-silent’ WM retention state and decoding the reactivated WM information, we demonstrate that content and structure are stored in a dissociative manner with distinct characteristics throughout WM process. First, each tone in the sequence is associated with two codes in parallel, characterizing its frequency and ordinal position, respectively. Second, during retention, a structural retrocue successfully reactivates structure but not content, whereas a following white noise triggers content but not structure. Third, structure representation remains stable whereas content code undergoes a dynamic transformation through memory progress. Finally, the noise-triggered content reactivations during retention correlate with subsequent WM behavior. Overall, our results support distinct content and structure representations in auditory WM and provide a novel approach to access the silently stored WM information in the human brain. The dissociation of content and structure could facilitate efficient memory formation via generalizing stable structure to new auditory contents.

## Introduction

Memories are not stored in fragments; instead, multiple items or events are constantly linked and organized with each other according to certain relationships. For a sequence of items to be successfully retained in working memory (WM), two basic formats of information need to be encoded and maintained in the brain – features that describe each item (content) and ordinal position of the item in the list (structure). For instance, making a phone call relies on correct assignment of ordinal labels to each digit, and rehearsing a piece of favorite melody requires sorting of each musical tone in certain temporal order. Thus, retaining even a simplified version of auditory temporally structured experience (e.g., a tone sequence) necessitates storage of two types of code – content (e.g., frequency for each tone) and structure (e.g., ordinal position for each tone) – in brain activities.

How does the human brain maintain content and structure information in WM system? Although amounts of previous studies have addressed the issue using different approaches in both animals and humans (e.g., Marshuetz, 2005; Komorowski et al., 2009; Eichenbaum, 2014; Kalm et al., 2014; Davachi & DuBrow, 2015; Kamiński & Rutishauser, 2020; Masse et al., 2020; Summerfield & Sheahan, 2020), most of the findings have been focused on the encoding period when the to-be-memorized stimuli are physically presented. Meanwhile, when entering the WM retention period, the brain would reside in a relatively ‘activity-silent’ state, during which information is ‘silently’ retained in synaptic weights of the network rather than showing sustained response to be explicitly decoded (e.g., Mongillo et al., 2008; Lewis-Peacock et al., 2012; Stokes, 2015, Rose et al., 2016, Masse et al., 2020). As a consequence, directly accessing neural representations of content and structure during retention and examining their WM behavioral relevance pose a huge challenge in the WM field.

Here in two auditory sequence WM experiments, we examined how content and structure information are encoded and maintained in auditory WM, by using a time-resolved multivariate decoding approach on the electroencephalography (EEG) recordings. Crucially, as mentioned previously, since the recorded macroscopic activities during the delay period tend to stay in an “activity-silent” state, it is difficult to directly and reliably assess the maintained WM information from EEG recordings. To this end, we employed an impulse-response approach (Wolff et al., 2017) to transiently perturb the neural network during retention and then measure the subsequently reactivated WM information. Specifically, two triggering events were presented successively during retention– a structure retrocue and a white-noise auditory impulse. The former, as an abstract structure cue, is hypothesized to reactivate the stored structure representation (i.e., ordinal position), while the latter – a neutral white-noise auditory impulse – might trigger content information (i.e., tone frequency) by perturbing the auditory cortex where contents likely reside.

Our results demonstrate that content and ordinal structure of an auditory tone sequence are encoded and maintained in a dissociative manner with distinct characteristics. First, each presented tone during encoding is associated with two codes in parallel, characterizing its frequency and ordinal position, respectively. Second, during the ‘activity-silent’ retention period, a structural retrocue successfully reactivates structure but not content, whereas a following white noise triggers content but not structure, implying their storage in different brain regions. Third, neural representation of structure information remains largely unchanged from encoding to retention whereas content code undergoes dynamical transformation, signifying their distinct representational formats. Finally and importantly, the white-noise-triggered content reactivations during retention correlate with subsequent memory performance, confirming its genuine indexing of WM operations. Taken together, our results provide new evidence advocating a dissociated and distinct form of content-structure storage in auditory WM and also constitute a novel approach to directly access WM information in human brains.

## Results

### Experimental procedure and time-resolved multivariate decoding analysis

In Experiment 1, thirty human participants performed an auditory delayed-match-to-sample working memory task while their 64-channel EEG activities were recorded. Each participant performed 1080 trials in total (6 hours in two separate days). As shown in Figure 1A, in each trial, three pure tones with different frequencies that were pseudo-randomly selected from six fixed values (*f*1: 381 *HZ, f*2: 538 *Hz, f*3: 762 *Hz, f*4: 1077 *Hz, f*5: 1524 *Hz, f*6: 2155 *Hz*) were presented sequentially, and participants were required to memorize both the ordinal position and frequency of the three tones. During the delay period, a retrocue (‘1’, ‘2’ or ‘3’) was first presented to indicate which of the three tones would be tested later, followed by a 100-ms white-noise auditory impulse. During the retrieval period, a target auditory tone was presented, and participants were instructed to determine whether its frequency was higher or lower than that of the cued tone. Note that the frequency of the target tone was determined in a pretest so that the overall memory performance would be around 75%. Indeed, the behavioral performance for the auditory WM task was 75.35% (SE = 1.00%).

**Figure 1.**
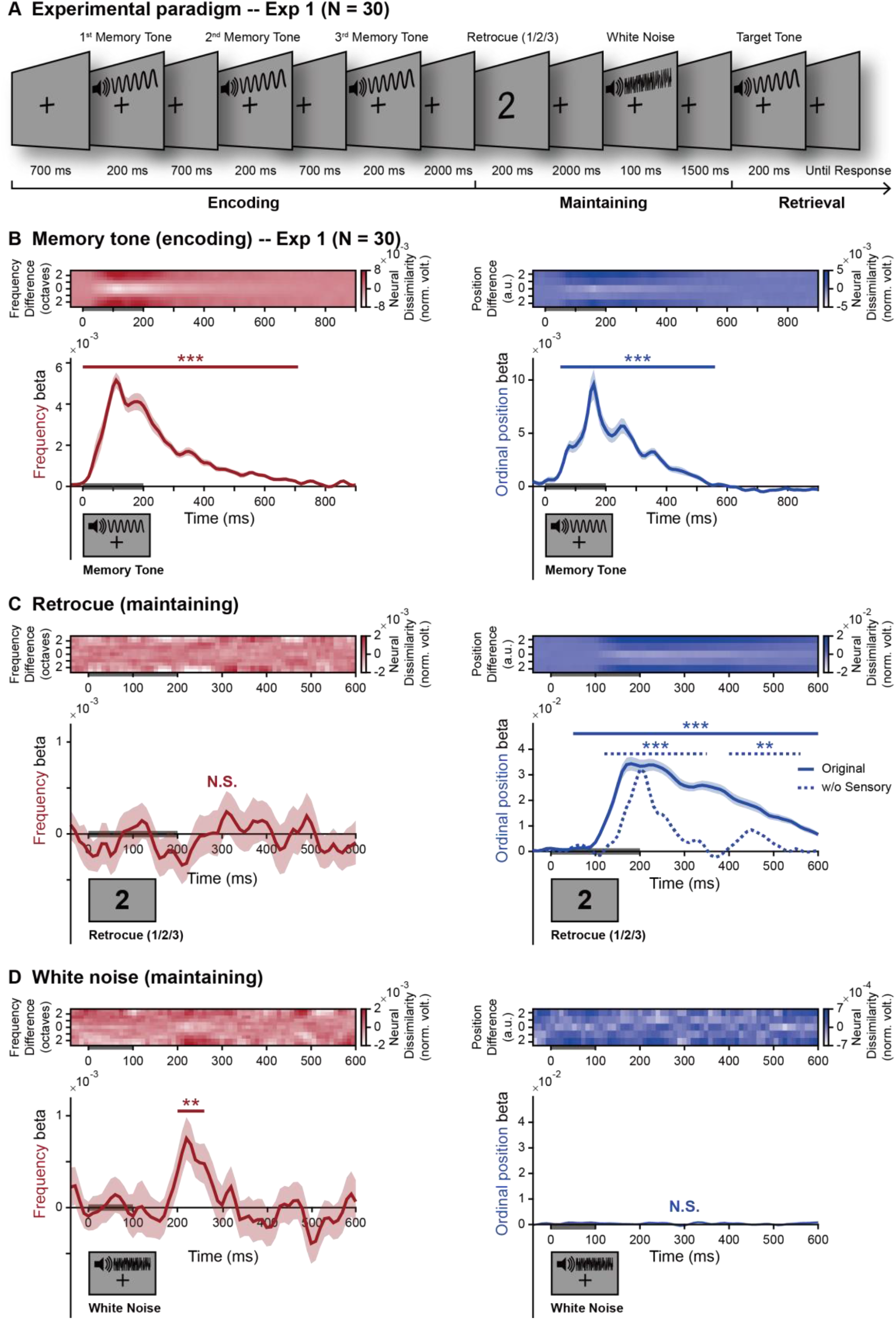
Experimental paradigm and neural representations of content and structure during encoding and retention periods (Exp 1) (A) Experimental paradigm (Exp1). In each trial, three pure tones (1^st^, 2^nd^, and 3^rd^ Memory Tones) with different frequencies (selected from 381 Hz, 538 Hz, 762 Hz, 1077 Hz, 1524 Hz and 2155 Hz) were serially presented and participants were instructed to memorize both frequencies and ordinal positions of the three tones. During the maintaining period, a retrocue (‘1’ or ‘2’ or ‘3’) appeared to indicate which of the three tones would be tested at the end of the trial. Next, a completely task-irrelevant white-noise auditory impulse was presented. During the retrieval period, a target pure tone was presented, and participants needed to compare it to the cued memory tone, i.e., higher or lower in frequency. (B) Neural representations of content (i.e., frequency) and structure (i.e., ordinal position) during encoding period. Upper panels: Grand average (N = 30) neural representational dissimilarity (mean-centered) as a function of physical dissimilarity of the memory tone, for frequency (left) and ordinal position (right), as a function of time. Lower panels: Grand average (N = 30, mean ± SEM) beta values of the regression between neural representational dissimilarity and physical dissimilarity of memory tones (i.e., decoding performance), for frequency (left) and ordinal position (right), as a function of time. Gray bar on x-axis indicates memory tone presentation. (C) Same as B but during retention period after retrocue. Gray bar on x-axis indicates retrocue presentation. Dotted blue line (right panel): sensory-information-removed decoding strength for ordinal position; corrected. (D) Same as B but during retention period after white-noise auditory impulse. Gray bar on x-axis indicates white-noise presentation. Note that only correct trials were analyzed in each participant. (Shaded area represents ± 1 SEM across participants. ^***^: p < 0.001; ^**^: p < 0.05; solid line: corrected using cluster-based permutation test, cluster-forming threshold p < 0.05).

A time-resolved representational similarity analysis (RSA) (Kriegeskorte et al., 2008) was performed on the EEG responses to evaluate the neural representation of frequency and ordinal position, respectively, throughout the encoding and maintaining phases, at each time point and in each participant (see details in Methods). Notably, instead of binary decoding, the RSA analysis here is based on a hypothesis that the neural representational similarity is proportional to the factor-of-interest similarity, i.e., the neural response for *f*1 (381 *Hz*) and *f*2 (538 *Hz*) would be less dissimilar than that for *f*1 (318 *Hz*) and *f*5 (1524 *Hz*). To quantify the representational strength, we next calculated the linear regression (*β*) between the factor dissimilarity and the corresponding neural representational dissimilarity for frequency and ordinal position, respectively. The decoding analysis was performed on content and structure information, respectively, i.e., decoding one factor (e.g., frequency) while marginalizing the other factor (e.g., ordinal position), and vice versa.

### Neural representations of structure and content during WM encoding

During the encoding period, each of the three tones could be characterized by two factors, i.e., frequency (*f*1, *f*2, *f*3, *f*4, *f*5, *f*6) and ordinal position (1^st^, 2^nd^, 3^rd^). Therefore, we could assess the neural representations of content and structure, respectively, from the same EEG response for each tone. Specifically, when examining frequency encoding, each tone would be labelled in terms of its frequency, regardless of its ordinal position, whereas analyzing structure representation would be based on ordinal positions regardless of the frequencies. Moreover, a cross-validated confound regression approach (Snoek et al., 2019) was employed to regress out possible influences from global field power courses for the three tones on the multivariate decoding analysis.

The upper panels of Figure 1B plot the time-resolved neural representational dissimilarity as a function of physical dissimilarity (y axis) after the onset of each tone, for frequency (left) and ordinal position (right). It is clear that for both properties, the neural dissimilarity is minimum at center and gradually increases to the two sides when the physical difference becomes larger, supporting that the neural responses contain information for both content and structure. The lower panel of Figure 1B shows the corresponding regression weights of the dissimilarity matrices, for frequency (*β*_*freq*_, left) and ordinal position (*β*_*pos*_, right), separately. Specifically, both the frequency representation *β*_*freq*_ (left; p < 0.001, corrected) and the ordinal position representation *β*_*pos*_ (right; p < 0.001, corrected) emerged shortly after the onset of each tone.

In summary, during the encoding period when a sequence of tones is presented to be retained in WM, each sound is signified by two separate neural codes that characterize its content (frequency) and structure (ordinal position), respectively (see Figure S1 for another analysis to confirm the separate encoding of frequency and ordinal position).

### Reactivations of structure and content information during ‘activity-silent’ retention

After establishing the representations of content and structure information during the encoding phase, we next examined their maintenance during retention when the brain responses enter an ‘activity-silent’ state (see Figure S2). Crucially, the triggering events – retrocue (Figure 1C) and white noise (Figure 1D) – successfully reactivated WM information from the silent state, yet content (left panel) and structure (right panel) displayed distinct temporal profiles. Specifically, as shown in Figure 1C, the retrocue triggered the emergence of ordinal position information (right, blue line) (p < 0.001, corrected) but not that for tone frequency (left, red line; p > 0.5, corrected). In contrast, the subsequent white-noise auditory impulse (Figure 1D) successfully activated the tone frequency representation (left, red line; 200 – 260 ms: p = 0.017, corrected) but not the ordinal position (right, blue line; p > 0.359, corrected). Thus, contents and ordinal structure are maintained in a dissociated way, i.e., being reactivated by different triggering events. Note that the RSA decoding analysis was performed in reference to the memory-related information, e.g., a trial with retrocue of ‘2’ and *f*4 (1077 *Hz*) as the 2^nd^ memorized tone frequency would be labelled as ‘2’ for ordinal position and *f*4 for frequency.

Importantly, since the retrocue stimuli are physically different (e.g., ‘1’, ‘2’, ‘3’), the observed structure representation after the retrocue (Figure 1C, right, blue solid line) would also contain the low-level sensory information. To address this issue, we further regressed the neural response dissimilarity according to two indexes – the ordinal positional dissimilarity (genuine structure information) and the dissimilarity between the neural response in the current experiment and that in a control experiment where participants viewed the identical retrocue stimuli (sensory information). This would allow us to remove the low-level sensory contributions from the structure decoding results. As shown in Figure 1C, the sensory-removed decoding performance remained present (right, dotted blue line), supporting that the ordinal position representations are not just caused by sensory inputs. It might still be argued that there are other confounding factors carried by the retrocue stimuli, an issue we further addressed using an encoding-to-maintaining generalization analysis (see Figure 2).

**Figure 2.**
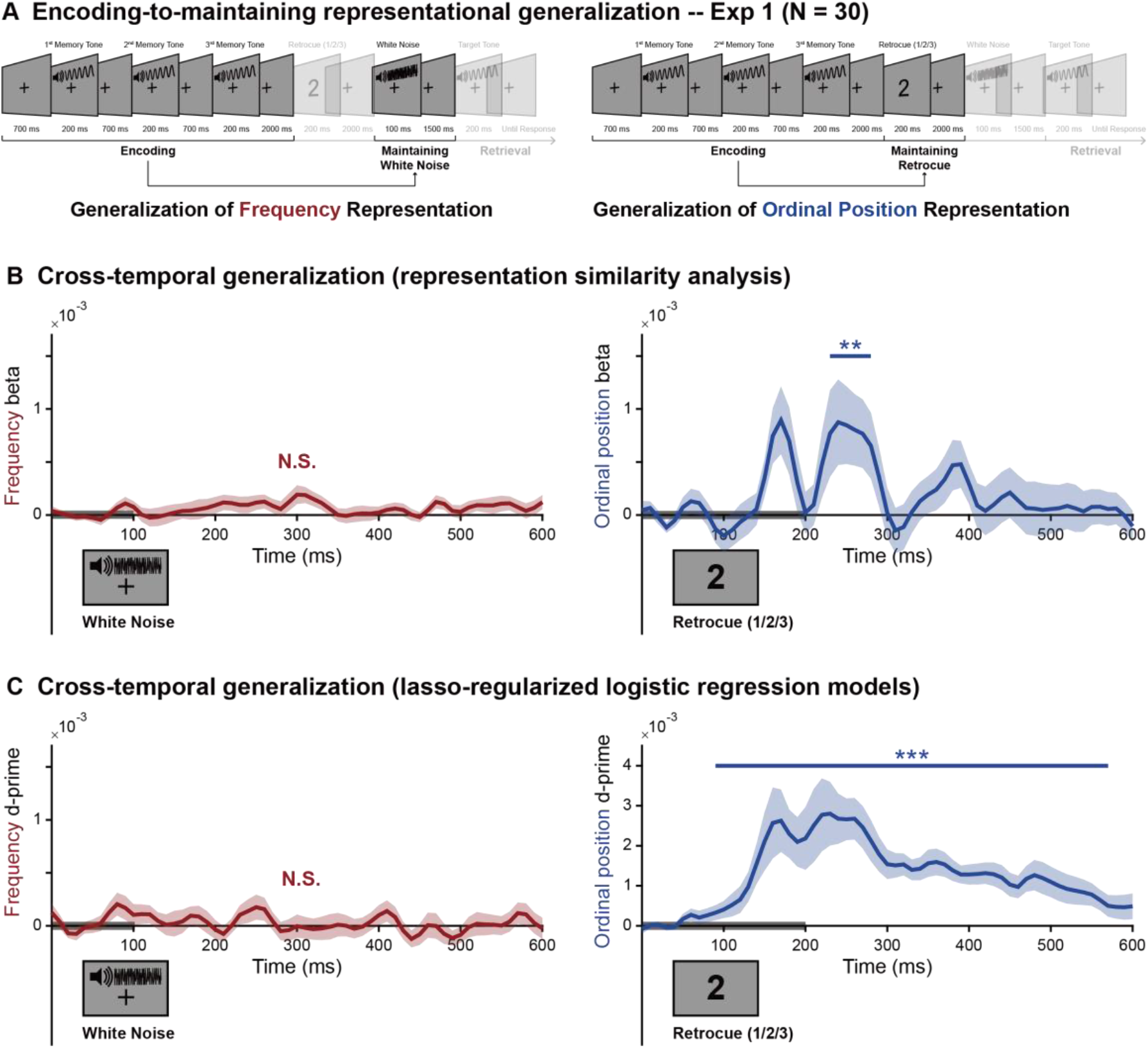
Encoding-to-Maintaining representational generalization (Exp 1) (A) Illustration of the cross-temporal generalization from encoding to maintaining periods for frequency (left) and ordinal position (right). Left: frequency generalization, i.e., encoding-phase response (within 90 - 130 ms) were used as train data to test the maintaining-phase responses at each time point after white-noise auditory impulse. Right: ordinal position generalization, i.e., encoding-phase response (within 140 - 180 ms) were used as train data to test the maintaining-phase responses at each time point after retrocue. (B) Grand average (N = 30, mean ± SEM) cross-temporal decoding generalization results using RSA as a function of time during maintaining, for frequency (left) and ordinal position (right). Gray bar on x-axis indicates white-noise (left) and retrocue (right). (C) Same as B but using lasso-regularized logistic regression models. Note that only correct trials were analyzed in each participant. (Shaded area represents ± 1 SEM across participants. ^***^: p < 0.001; ^**^: p < 0.05; solid line: corrected using cluster-based permutation test, cluster-forming threshold p < 0.05).

Another concern is whether the white noise impulse presented during retention would interfere with auditory WM. A control experiment with completely different participants (N =19; see Figure S3) excluded the possibility, by revealing similar WM behavioral performance with or without noise. The results further support that the white noise serves as a ‘neutral’ probe to assess the information retained in WM network.

Overall, during the ‘activity-silent’ maintaining period, a structural retrocue successfully reactivates structure but not content information, whereas a following white-noise auditory impulse triggers WM content (to-be-recalled tone frequency) but not structure representation, implying their storage in different brain regions.

### Distinct encoding-to-maintaining representational generalizations for structure and content

Having confirmed the neural representations during both encoding (Figure 1A) and maintaining periods for structure (Figure 1C, after retrocue) and content (Figure 1D, after white noise), we next conducted an encoding-to-maintaining generalization analysis to estimate whether these two phases share similar coding formats. Put another way, if the neural codes for encoding (Figure 1B) could be successfully generalized to those elicited during retention (Figure 1CD), this would support a stable WM coding, otherwise a dynamic coding transformation.

We first used the same RSA approach as before to examine the cross-temporal generalization, except that here the response dissimilarity was calculated between the encoding and retention activations. As shown in Figure 2B, only the structural information showed a significant cross-temporal generalization (right; 230 – 280 ms: p = 0.026, corrected), whereas the content information did not (left; p > 0.089, corrected). To further verify the results, we used a lasso-regularized logistic regression models to test the cross-temporal generalization. As shown in Figure 2C, likewise, only the structure information showed significant encoding-to-maintenance generalization (right; 90 – 570 ms: p < 0.001, corrected), but not for frequency information (left; p > 0.438, corrected). Importantly, the successful cross-temporal generalization for ordinal position further confirmed the genuine structure information triggered by the retrocue during retention, since no visual cues (i.e., ‘1’, ‘2’, ‘3’ stimuli) were presented during the encoding period yet the structural codes could still be generalized to that during retention.

Taken together, structure and content are endowed with distinct representational transformation properties, i.e., structure representation remains stable from encoding to maintaining periods, while the content code undergoes a dynamic transformation, signifying their distinct representational formats.

### Content reactivations during retention correlate with WM behavior

Finally, we evaluated whether the neural representations of content and structure retained in WM, which could be reactivated by the retrocue and white noise respectively, have any behavioral consequence. To this end, we performed the same RSA analysis on correct trials and incorrect trials separately, in each participant. We ensured equal numbers of correct and incorrect trials to make a fair comparison (see details in Methods). Interestingly, as shown in Figure 3A, the correct (dark red) and wrong trials (light red) displayed distinct frequency reactivations after the white noise (correct vs. wrong: black asterisks; 210 – 250 ms: p = 0.053, corrected). Specifically, the correct-trial group (dark red) showed significant frequency decoding performance (200 – 260 ms: p = 0.018, corrected), whereas the wrong-trial group (light red) did not. Interestingly, the wrong-trial group also showed a trend of late reactivation (440 – 460 ms: p = 0.178, corrected; maximum p = 0.049, uncorrected), with a trend of behavioral relevance (wrong vs. correct: 440 – 450 ms: p = 0.351, corrected; maximum p = 0.016, uncorrected).

**Figure 3.**
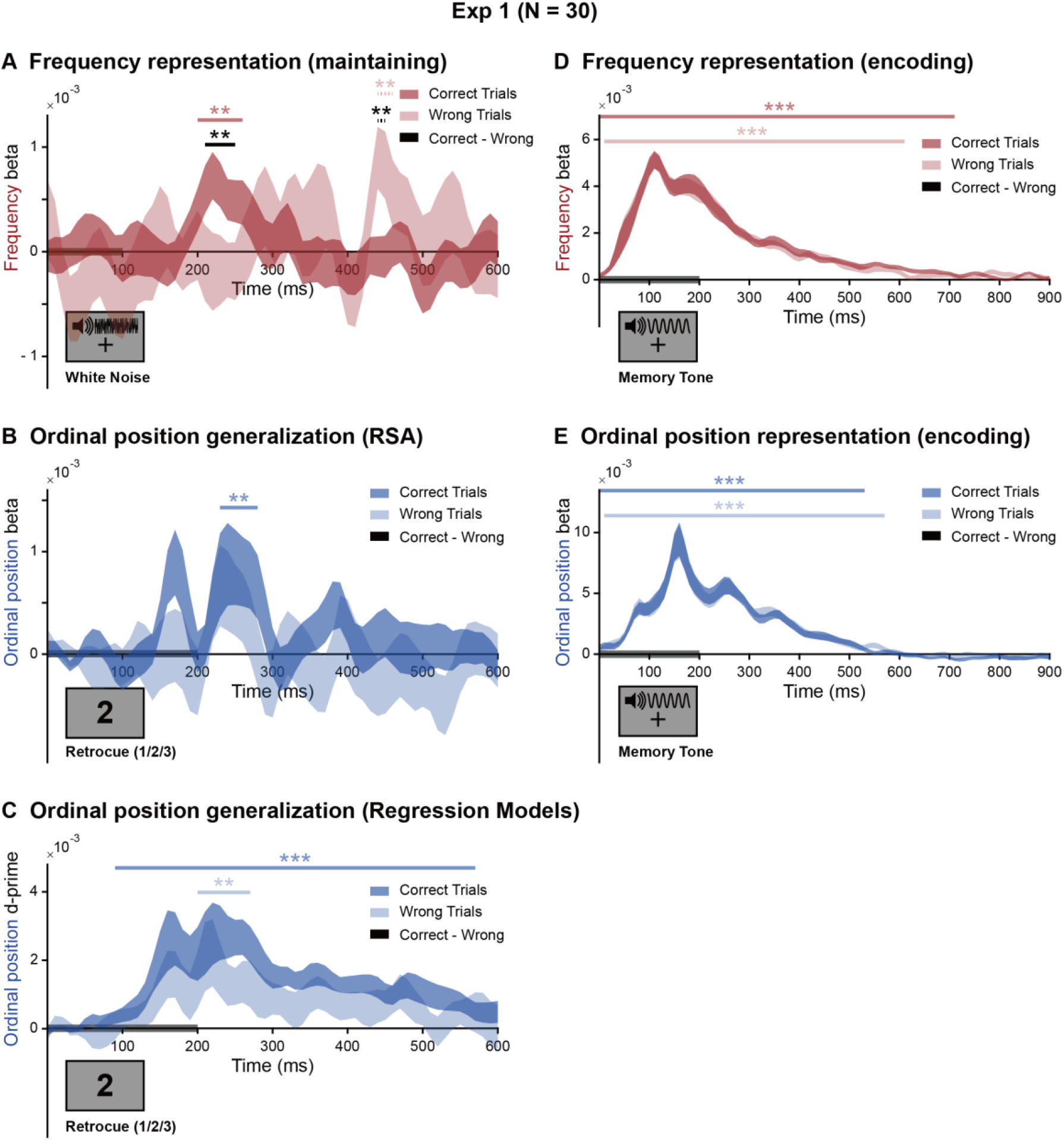
Behavioral correlates of content and structure representations in auditory WM (Exp 1) (A) Grand average (N = 30, mean ± SEM) frequency decoding performance during maintaining period after white-noise auditory impulse, for correct (dark red) and wrong (light red) trials. Gray bar on x-axis indicates white-noise presentation. (B) Grand average (N = 30, mean ± SEM) structure decoding performance during maintaining period after retrocue (i.e., encoding-to-maintaining generalization), for correct (dark blue) and wrong (light blue) trials. Gray bar on x-axis indicates retrocue presentation. (C) Same as B but using lasso-regularized logistic regression models. (D) Same as A but during encoding period. Gray bar on x-axis indicates memory tone presentation. (D) Same as B but during encoding period. Note that the decoding analyses were performed on correct and wrong trials with equal trial numbers. (Shaded area represents ± 1 SEM across participants. ^***^: p < 0.001; ^**^: p < 0.05; solid line: corrected using cluster-based permutation test, cluster-forming threshold p < 0.05; dotted line: uncorrected).

Meanwhile, the structure reactivations (i.e., encoding-to-maintaining generalization, given its genuine indexing of structure information independent of retrocue stimulus) displayed no behavioral relevance, i.e., correct and wrong trials showed comparable structure reactivations (RSA, Figure 3B, p > 0.138, corrected; Lasso-regularized logistic regression model, Figure 3C, p > 0.106, corrected). Furthermore, the neural representation during the encoding period did not show behavioral correlates for both frequency (Figure 3D, p > 0.287, corrected) and ordinal position (Figure 3E, p > 0.5, corrected), suggesting that the deteriorating WM performance in wrong trials were not attributable to encoding failure.

Thus, content information (i.e. tone frequency) maintained in WM, which could be assessed using a neutral white-noise auditory impulse during retention, covaries with subsequent WM recalling performance, thus further confirming the reactivation’s genuine indexing of WM operations.

### Experiment 2: task control

In Experiment 1, participants were instructed to retain a list of auditory tones and recall the tone frequency according to the retrocue (1^st^, 2^nd^, 3^rd^). Meanwhile, since the recalling task only tested the content, participants might have discarded the structure information after the retrocue. This would constitute an alternative interpretation for why the white-noise impulse failed to trigger structure representations (Figure 1CD). To address this possibility, we performed Experiment 2 (N = 18), during which the participants did exactly the same task as Experiment 1 but needed to additionally recall the position of the memorized tone (Figure S4A).

The behavioral accuracies for frequency and ordinal position were 75.17% (SE = 1.08%) and 98.80% (SE = 0.22%), respectively. As shown in Figure 4, Experiment 2 largely replicated the results of Experiment 1 (see Figure S4B for results during encoding period in Exp 2), thus excluding the task demand interpretation. Specifically, the retrocue elicited structure representation (Figure 4A, right; 60 – 600 ms: p = 0.003, corrected) but not frequency (Figure 4A, left; p > 0.337, corrected), whereas the white noise reactivated content (Figure 4B, left, dark red; 250 – 320 ms: p = 0.006, corrected) but still failed to drive structure information (Figure 4B, right, p > 0.350, corrected). Moreover, same as Experiment 1, the cross-temporal generalization results showed a significant encoding-to-maintaining generalization for structure (Figure 4C, right, dark blue; 140 – 180 ms: p = 0.046, corrected) but not for content (Figure 4C, left, p > 0.251, corrected). Finally, also consistent with Experiment 1, correct trials showed better content reactivations than wrong trials (Figure 4B, left, black star, 250 – 280 ms: p = 0.054, corrected), but not for ordinal position (Figure 4C, right p > 0.158, corrected).

**Figure 4.**
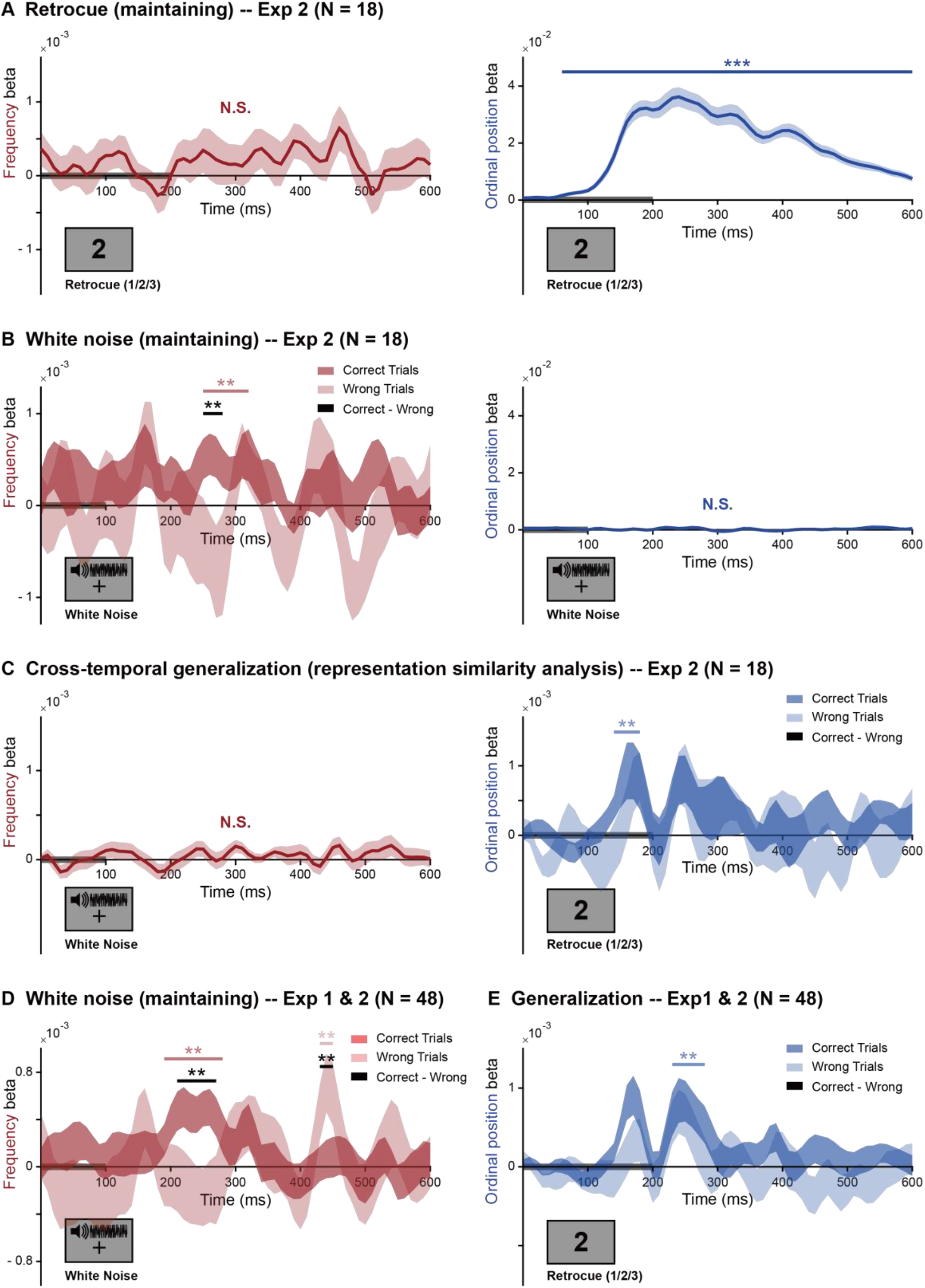
Experiment 2 results. (A) Neural representations of content (i.e., frequency) and structure (i.e., ordinal position) during maintaining period after retrocue. Grand average (N = 18, mean ± SEM) beta values of the regression between neural representational dissimilarity and physical dissimilarity of cued tones, for frequency (left) and ordinal position (right), as a function of time. (B) Left: Grand average (N = 18, mean ± SEM) frequency decoding performance after white noise, for correct (dark red) and wrong (light red) trials. Right: Grand average ordinal position decoding performance after white noise. (C) Grand average (N = 18, mean ± SEM) encoding-to-maintaining generalization results for frequency (left) and ordinal position (right) for correct (dark blue) and wrong (light blue) trials. (D) Grand average (N = 48, mean ± SEM) frequency decoding performance after white noise, for correct (dark red) and wrong (light red) trials, by combining Exp 1 and Exp 2. (E) Grand average (N = 48, mean ± SEM) encoding-to-maintaining generalization results for ordinal position after retrocue, for correct (dark blue) and wrong (light blue) trials, by combining Exp 1 and Exp 2. Note that the analyses were performed on correct and wrong trials with equal numbers in each participant. (Shaded area represents ± 1 SEM across participants. ^***^: p < 0.001; ^**^: p < 0.05; solid line: corrected using cluster-based permutation test, cluster-forming threshold p < 0.05; dotted line: uncorrected).

We next combined Experiment 1 and Experiment 2 (N = 48), given that they shared the same design through encoding and retention periods and showed similar activation patterns. Consistently, the white noise triggered stronger content reactivation for correct than incorrect trials (Figure 4D, black, 210 – 270 ms: p = 0.011, corrected), while the retrocue did not (Figure 4E, p > 0.148, corrected). Intriguingly, there was still a trend of late content reactivation for incorrect trials (430 – 450 ms: p = 0.179, corrected; maximum p = 0.045, uncorrected), which was even marginally higher than that for correct trials (430 – 450 ms: p = 0.178, corrected; maximum p = 0.045, uncorrected), implying that the WM content in wrong trials might not be lost completely, but tends to be maintained in a less excitable latent state of WM network (see Discussion).

Taken together, the new control experiment fully replicated previous results, supporting that the lack of structure reactivation after the white-noise impulse is not due to task demands in Experiment 1.

## Discussion

As we learn about the world, memories are formed not only for individual events but also for their relational structure. Here, we demonstrate that content (tone frequency) and structure (ordinal position) of a tone sequence are encoded and stored in a distinct way in human auditory WM. Each item is signified by two separate neural codes – content and structure – which could be subsequently reactivated from the ‘activity-silent’ state during WM retention, by white-noise auditory impulse and retrocue, respectively. Furthermore, content and structure demonstrate distinct temporal properties throughout memory process, i.e., stable representation for structure and dynamical coding for content. Importantly, the content reactivations during retention are further correlated with WM recalling performance. Overall, the human brain extracts and separates structure and content information from auditory inputs and maintains them in distinct formats in the WM network. Our work also provides a novel approach to access the ‘silently’ stored WM information in the human brain.

It has long been posited that structure information that characterizes the abstract relationship between objects serves as a primary rule to organize fragmented contents and influences perception (i.e., Gestalt principle; Wagemans et al., 2012), memory (Davachi & DuBrow, 2015; DuBrow & Davachi, 2013; Gershman et al., 2013) and learning (Luyckx et al., 2019). How is the structure information implemented in the brain? One typical example is the hippocampus neurons that are sensitive to specific spatial location or time bin that are independent of the attached contents, signifying their coding of spatial (Eichenbaum et al., 1999; Muller, 1996; O’Keefe & Dostrovsky, 1971; O’keefe & Nadel, 1978), temporal (Eichenbaum, 2014; MacDonald et al., 2011; Manns et al., 2007; Paz et al., 2010; Tsao et al., 2018), and abstract structures even in nonspatial tasks, i.e., cognitive map (Aronov et al., 2017; Garvert et al., 2017; Schuck & Niv, 2019). Parietal regions are also involved in structural representation and learning (Feigenson et al., 2004; Summerfield et al., 2020;). Here we focused on the ordinal information of a tone sequence, a typical and important relational structure to sort contents in auditory experience. Indeed, neural coding of ordinal position has been found in both animal recordings (Fortin et al., 2002; Naya & Suzuki, 2011) and neuroimaging studies (Baldassano et al., 2017; Carpenter et al., 2018; DuBrow & Davachi, 2016; Hsieh et al., 2014; Kikumoto & Mayr, 2018; Liu et al., 2020; Rajji et al., 2017; Roberts et al., 2013; Huang et al., 2018). Our results are thus consistent with previous findings and further expand to auditory sequence memory, particularly during the maintaining period.

Different from visual studies, only a few studies have assessed the neural correlates of auditory WM (Albouy et al., 2017; Kumar et al., 2016; Luo et al., 2013; Weisz et al., 2020; Wolff et al., 2020). Here, by combining RSA-based decoding and impulse-response approach, we demonstrate that a neutral white-noise auditory impulse could successfully reactivate the tone frequency information ‘silently’ held in auditory WM. This is consistent with the hidden-state WM view, which posits that information is stored in synaptic weights via short-term neural plasticity principles (Lewis-Peacock et al., 2012; Masse et al., 2020; Mongillo et al., 2008; Stokes, 2015). As a result, a transient disturbance of the neural network (e.g., a flash or a white noise) would endow the neural assemblies that maintain WM information in synaptic weights with larger probability to be activated (Stokes, 2015). In other words, the white-noise here serves as a ‘neutral’ probe to assess the information that has already been retained in WM and would not modify WM contents (also see the control experiment, Figure S3). Nevertheless, not all the triggering events could efficiently reactivate contents, e.g., the retrocue failed to trigger content information, implying that content and structure might be stored in different regions and have different characteristics. Auditory content, presumably retained in sensory cortex, is sensitive to auditory perturbation, while structure information might be maintained in high-level areas, e.g., parietal region (Bueti et al., 2009; Parkinson et al., 2014; Summerfield et al., 2020) and frontal cortex (Berdyyeva & Olson, 2010; Hsieh et al., 2011; Naya et al., 2017; Ninokura et al., 2003).

The white-noise-elicited content reactivation during retention correlates with subsequent recalling performance, further supporting its genuine indexing of content maintenance in auditory WM. Interestingly, a trend of late content reactivation for incorrect trials suggests that content information might not be completely lost but still retained in WM, yet in a less excitable latent state, i.e., being triggered in a late and less robust way. A recent visual WM study reveals that top-down attention could modulate the latent states of WM items, i.e., the most task-relevant one would be in a more excitable state and tends to be reactivated earlier by TMS perturbation (Rose et al., 2016). Moreover, computational modelling that incorporates competition between items also predicts late reactivation for weakly stored WM items (Mongillo et al., 2008). Therefore, besides reactivation strength, the latency of content reactivations might also serve as a potential index to assess subsequent WM performance.

What would be the benefit of the dissociated content and structure representation in WM system? An apparent advantage is that this would allow rapid and versatile transfer of the structure information to new contents (Behrens et al., 2018; Friston et al., 2016; Kaplan et al., 2017; Tse et al., 2007; Whittington et al., 2018). Visual WM studies also reveal that different attributes (e.g., color, shape or orientation) of a single object are coded independently (Bays et al., 2011; Cowan et al., 2013; Fougnie & Alvarez, 2011). Back to the current experiment, all trials share the fixed sequence structure (e.g., 3-tone sequence), and it would therefore be more efficient for the brain to separately encode and maintain the ordinal position information and allocate tone information anew in each trial to the corresponding ordinal positions.

Finally, structure and content also show different coding dynamics, i.e., stable and dynamic representations, respectively. The transformed code for WM contents has been widely found (e.g., Barak et al., 2010; Kamiński & Rutishauser, 2020; Lundqvist et al., 2016, 2018; Meyers et al., 2008; Parthasarathy et al., 2017, 2019; Quentin et al., 2019; Rademaker et al., 2019; Spaak et al., 2017; Sprague et al., 2016; Trübutschek et al., 2017; Wolff et al., 2017, 2020; Yu et al., 2020; Yue et al., 2019). Possible interpretations for the dynamic coding are that WM content is coded in a way that is optimized for subsequent behavioral demands (Panichello & Buschman, 2020; Myers et al., 2017), or transformed into a subspace to resist distractions from new inputs (Libby & Buschman, 2019; Murray et al., 2017). In contrast, the ordinal structure displays a stable representation over memory course, in line with previous findings (Kalm & Norris, 2017; Luyckx et al., 2019; Walsh, 2003). Stable structure coding would be advantageous for memory generalization and formation, i.e., rapid implementation of stable structure to dynamic contents.

Taken together, content (tone frequency) and structure (ordinal position) are the two basic formats of information to be maintained in WM. Our findings demonstrate that they are encoded and stored in a largely dissociated way and display distinct representational characteristics, which echo their varied functions in memory formation – detailed and dynamic contents vs. abstract and stable structure.

## Author contributions

Y.F. and H.L. designed the experiment. Y.F. and S.G. performed the experiment. Y.F. and Q.H. analyzed the data. Y.F. and H.L. wrote the paper.

## Acknowledgments

This work was supported by the National Natural Science Foundation of China (31930052 to H.L.), Beijing Municipal Science & Technology Commission (Z181100001518002) to H.L.

## METHODS

### Participants

Thirty (17 females, mean age 23.4, range 19-27 years) and twenty (8 females, mean age 19.7, range 18–25 years) healthy participants with normal or corrected-to-normal vision were recruited in Experiment 1 and Experiment 2, respectively, after providing written informed consent. Two participants in Experiment 2 were excluded due to low behavioral performance. Participants received compensation for participation. The experiment was approved by the Departmental Ethical Committee of Peking University.

### Apparatus and Stimuli

The experiment stimuli were generated and controlled with MATLAB (MathWorks) and Psychophysics Toolbox (Psychtoolbox-3; Brainard, 1997). The visual stimuli were presented on a Display ++ LCD screen with a resolution of 1920 by 1080 pixels running at a refresh rate of 120 Hz. The distance between the screen and participants was fixed at 60 cm. Auditory stimuli were presented with a Sennheiser CX213 earphone through an RME Babyface pro external sound card. The intensity of all auditory stimuli was set at approximately 65 dB (62.1 to 67.3 dB) SPL. A USB keyboard was used for response collection.

### Experimental procedure

#### Experiment 1

Each trial started with the presentation of a cross (0.9° visual angle), which stayed in the center of the screen throughout the entire trial except during the retrocue’s presentation (Figure 1A). Participants were instructed to fixate at the central cross during the entire trial. After 700 ms, three memory tones with different frequencies that are pseudo randomly selected from a uniform distribution of 6 frequencies (381 Hz, 538 Hz, 762 Hz, 1077 Hz, 1524 Hz and 2155 Hz) were presented sequentially. The duration of each memory tone was 200 ms, with inter-stimulus interval of 700 ms. 2000 ms after the offset of the third tone, a 200-ms visual retrocue (1.9° visual angle), a visual character (‘1’, ‘2’ or ‘3’), was presented in the center of the screen, indicating which of the three memory tones would be tested later, with 100% validity. A 100-ms white-noise auditory impulse was then presented 2000 ms after the retrocue. Finally, after another 1500-ms interval, a 200-ms target auditory tone was presented. Participants made a judgement on whether the frequency of the target tone was higher or lower than the frequency of the cued memory tone, by pressing keys on the keyboard, without time limitation. The frequency of the target tone was either 2^1/3^ higher or lower, with equal probability, than the frequency of the cued memory tone. Next trial started with a 1500 - 2000 ms delay after participant made their responses. There were 1080 trials in total, which were separated into two sessions that were completed in two separate days, for each participant. Each session lasted approximately 3 hours.

#### Experiment 2

Experiment 2 was the same as Experiment 1, except that participants were instructed to additionally report the ordinal position of the cued tone during recalling (see Figure S4A). Furthermore, to control the temporal length of the experiment, there were other timing parameter adjustments, i.e., the interval between the 3^rd^ auditory tone and the retrocue was set to 1500 ms, the interval between the retrocue and white noise to 1700 ms, and the interval between white noise and target tone to 1000 ms. Moreover, in order to prevent motor preparation during the maintaining period, the correspondence between particular ordinal positions and reaction keys are randomly set from six combinations in each trail. Same as Experiment 1, there were 1080 trials in total, which were separated into two sessions being completed in two separate days, for each participant. Each session lasted approximately 3 hours.

### EEG acquisition and pre-processing

The EEG signals were acquired using a 64-channel actiCAP (Brain Products) and two BrainAmp amplifiers (Brain Products). The data was recorded through BrainVision Recorder software (Brain Products) at 500 Hz. Vertical electrooculogram was recorded by one additional electrode below the right eye. The impedances of all electrodes were kept below 10 kΩ. The EEG data was referenced to the average value across all channels, down-sampled to 100 Hz, and bandpass filtered between 1 and 30 Hz. Independent component analysis was performed to remove eye-movement and other artifactual components. Data were epoched - 500 ms before each trial’s onset to 700 ms after target tone’s offset. Epochs with extremely high noises by visual inspection were manually excluded from the following analyses.

### Time-resolved multivariate decoding

Note that only the correct trials in each participant were used for further analysis, except the behavioral correlates analyses (Figure 3, Figure 4) during which the same number of correct and wrong trials in each participant were analyzed and compared (see details in later Methods).

First, a time-resolved representational similarity analysis (RSA) was performed on the EEG data to evaluate the neural representation of frequency and ordinal position independently throughout the encoding and maintaining phases, at each time point for each participant. To remove the possible interference of slow trend, we first calculated the mean activity for each channel across all trials, and then the trial mean was smoothed by a 150-ms moving window and subtracted from the original data trial wisely (i.e., demean; Grootswagers et al., 2017). In addition, as the three pure tones were presented successively during encoding period, the difference of global field power (GFP) of these three tones might contribute to the ordinal position decoding analysis. To remove the possible confounding effects, we employed a cross-validated confound regression method proposed by Snoek et al. (2019) to remove the variance that could be explained by the GFP levels from the encoding responses. All the subsequent decoding analysis were conducted on the corrected data.

The RSA analysis is based on both spatial and temporal information to achieve high signal-to-noise ratio (Grootswagers et al., 2017). Specifically, for RSA-based decoding at time point *t*, the values of all channels at the current time point *t* as well as those at previous time point *t-1* were all included as features (64^*^2 = 128 in total) for a further 8-fold cross-validation decoding approach. The Mahalanabis distance (Mahalanobis, 1936) in the neural spatiotemporal activities (i.e., 128 features) between each left-out test trial and the averaged, condition-specific activities over train-trials was computed, with the covariance matrix estimated from all the train-trials using a shrinkage estimator (Ledoit & Wolf, 2004). If the neural activities indeed contain information about certain feature (e.g., tone frequency, ordinal position), the more similar the two features are (less physical dissimilarity), the less Mahalanobis distance their associated neural activities would have (less neural representational dissimilarity). To further quantify their relationships, the physical differences were linearly regressed against the corresponding neural representational dissimilarity values, for each test-trial. The mean beta value of the regression across all test-trials was then used to represent the decoding accuracy. This process was repeated 50 times with each containing a new random partition of data into 8 folds. The resulted decoding performance, smoothed with a Gaussian-weighted window (window length = 40 ms), were then averaged across the 50 partitions as the final decoding accuracy, at each time point and for each participant. Note that the RSA analysis was performed for frequency and ordinal position, independently, using the same EEG data but with different feature dimensions.

Notably, the ordinal position decoding results would contain the low-level sensory information carried by the retrocue stimuli (i.e., ‘1’, ‘2’, ‘3’). To remove the confounding influences, we ran a control experiment during which participants were instructed to passively view the same retrocue stimuli (‘1’, ‘2’, ‘3’) and their EEG responses were recorded. We then conducted a leave-one-participant-out decoding analysis on our main experiment data. Specifically, in each run, the response of one participant in the main experiment was used as test data, with the remaining participants’ responses in the main experiment as train dataset 1 and those in the control experiment as train dataset 2. The neural response representational dissimilarity between the test data and train dataset 1 (containing both structural and sensory representations), and that between the test data and train dataset 2 (containing only sensory representations) were then calculated, respectively. Finally, the neural dissimilarity between the test data and train dataset 1 (containing both structural and sensory representations) was regressed according to two predictors – the ordinal positional dissimilarity (structure information) and the sensory neural dissimilarity (calculated between test data and train dataset 2) – with the former referring to the sensory-removed representational strength (dotted line in Figure 1C, right).

For correct and wrong trial decoding analyses, the right trials were k-folded, where k was determined by the closest integer to the quotient of the total correct trial numbers and wrong trial numbers. For each partition, the (k-1)-folds correct trials were used as training data, and the left 1-fold correct trials and all wrong trials were used as testing data, respectively. This manipulation will ensure the same trial numbers for correct and wrong trials during testing. The above k-fold decoding procedure was further conducted 50 times, and averaged to obtain the final decoding time courses.

### Cross-temporal generalization analysis

A cross-temporal generalization analysis was conducted to investigate whether the neural representations of frequency or ordinal position were similar for encoding and maintaining periods. We first selected the temporal range during the encoding period to serve as that for training data, based on their corresponding decoding performance during encoding (i.e., 90 - 130 ms and 140 - 180 ms after the onset of memory tone, for frequency and ordinal position, respectively). Note that as shown in Figure 1CD, the neural representations of frequency and ordinal position only appeared after the white-noise stimulus and retrocue, respectively, during the delay period. Therefore, the cross-temporal generalization was performed only on the corresponding time ranges for these two features, i.e., 0 – 600 ms in reference to the onset of white-noise stimulus and retrocue, for frequency and ordinal position, respectively.

We employed two types of multivariate decoding methods – RSA and lasso-regularized logistic regression model – for the cross-temporal generalization analysis. RSA was similar to previous analysis, except that the Mahalanibos distance was computed between responses during the encoding period (train data) and that during the maintaining period (test data). For lasso-regularized logistic regression model, we first trained classifier 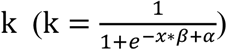, with *x* as 128 decoding features (64 channels over 2 time points) derived from EEG responses, vector *β* representing weights for each feature and *α* as the intercept coefficient, to differentiate either ordinal positions or frequencies using the encoding-phase responses. The obtained classifiers were then applied on the maintaining-phase data. The output was next transformed to d-prime values, which represent the cross-temporal generalization decoding performance.

For both methods, the decoding results were further baseline corrected (0 to 50 ms), repeated for 50 times, and then averaged. Finally, to remove possible influences from random effects, a control analysis was performed by shuffling test data labels and redoing the same cross-temporal generalization analysis for 50 times. The resulted shuffling results were then subtracted from the original results to get the final cross-temporal generalization results.

### Statistical significance testing

Non-parametric sign-permutation test (Maris & Oostenveld, 2007; customized analysis codes) was used for statistical test. Specifically, the sign of the decoding value of each participant at each time point was randomly flipped 100,000 times to obtain the null distribution, from which the p value was derived. Cluster-based permutation test was then conducted to correct multiple comparisons over time (p < 0.05). For clusters failing to pass the multiple comparison, the uncorrected results were reported.

