## Supplementary Figure 1-4 for "Distinct neural representations of content and ordinal structure in auditory sequence memory"

### Supplemental Information

#### Figure S1

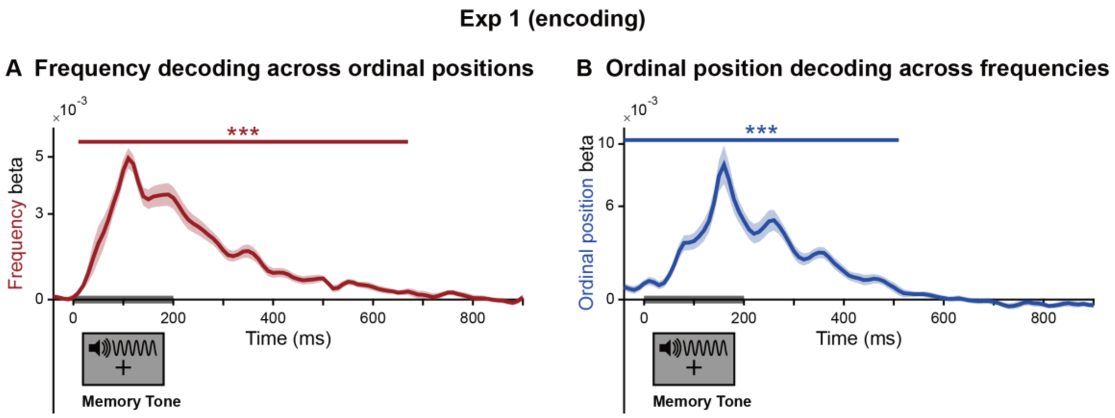


**Figure S1. Control analysis during encoding period (Exp 1, N =30).**
For frequency decoding, we constrained the training data on one specific position (e.g., the 1$^{st}$ position) and tested the frequency decoding on data at the 2$^{nd}$ and 3$^{rd}$ position, respectively. Likewise, for ordinal position information, we constrained the training data on one specific frequency (e.g., 538 Hz) and tested the position decoding on data for the other 5 frequencies, respectively. Both frequency (A) and ordinal position (B) showed robust decoding generalization across other positions and other frequencies, respectively (Shaded area represents ± 1 SEM across participants. ***: $p < 0.001$; **: $p < 0.05$; solid line: corrected using cluster-based permutation test, cluster-forming threshold $p < 0.05$).

#### Figure S2

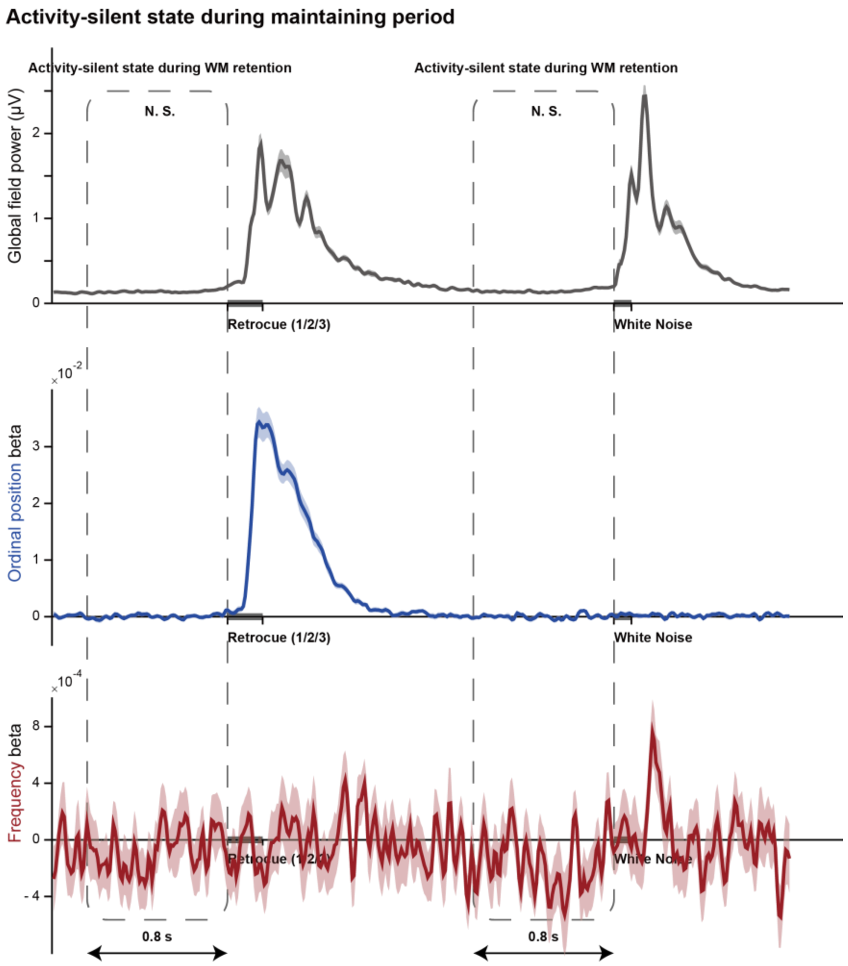


**Figure S2. Activity-silent state during WM retention before retrocue or white noise (Exp 1, N =30).**

The full-range data throughout the WM retention period for global field power (upper panel), ordinal position decoding (middle panel), and frequency decoding (lower panel). It is clear that neither global field power nor multivariate results show activations before the retrocue and white noise during the delay period (dotted box), supporting the ‘activity-silent’ view. (Shaded area represents ± 1 SEM across participants.)

#### Figure S3

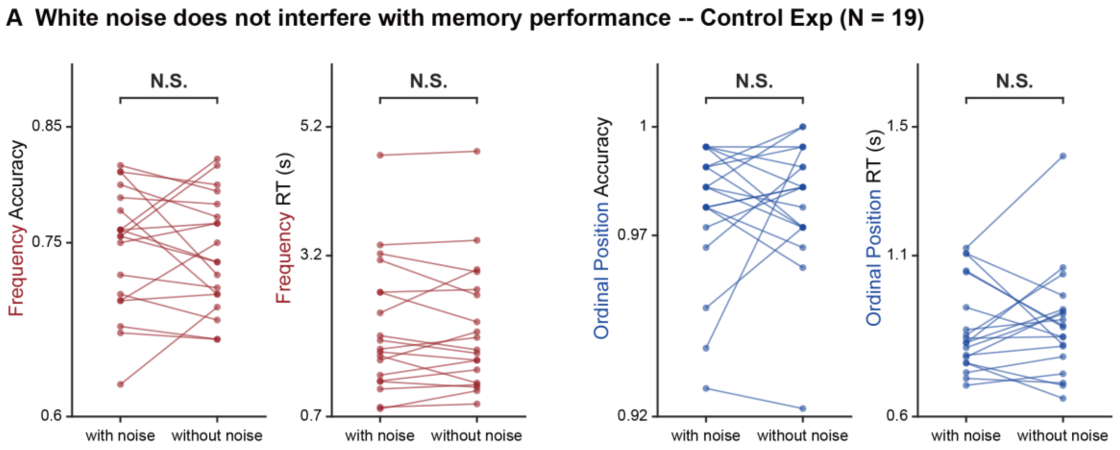


**Figure S3. Control behavioral experiment (N = 19).**

This behavioral experiment (N = 19, 12 females; different participant groups from Exp 2) is to examine whether the added white noise would interfere with auditory WM. Participants did the same WM task (Figure S4 A) as in Exp 2, except with or without presenting the white noise during the delay period. There were 180 trials for each condition (360 trials totally). Each dot represents individual result. The two conditions (with noise, without noise) showed no behavioral difference in both accuracy and RT on content and structure memory, thus supporting that the presented white noise during retention does not interfere with auditory WM.

#### Figure S4

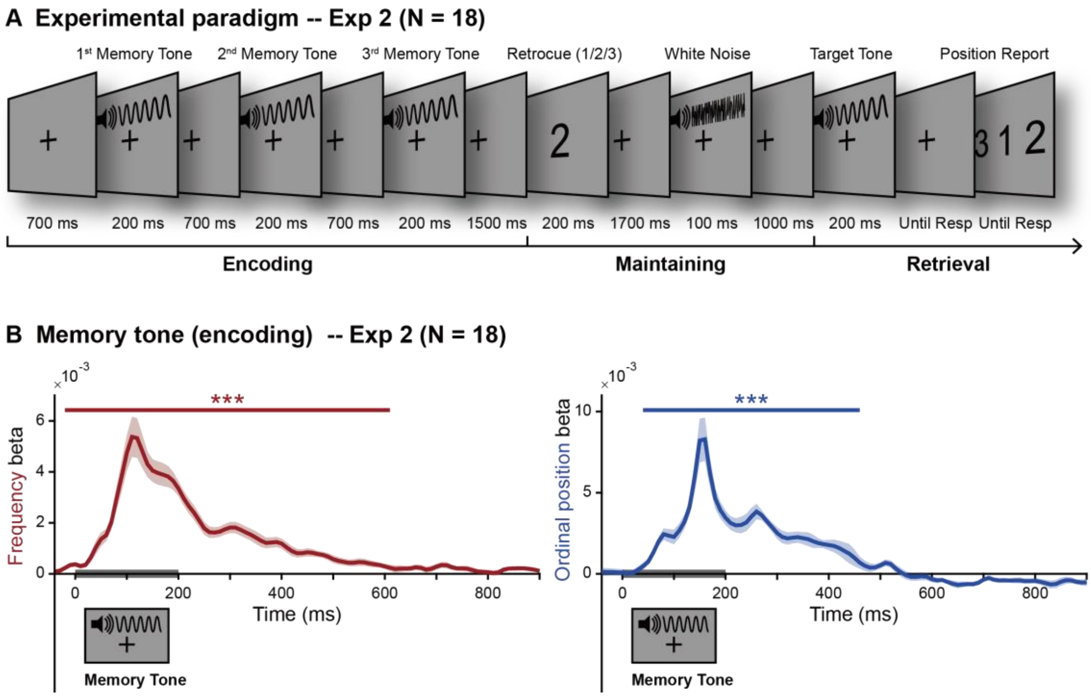


**Figure S4. Experiment 2 paradigm and the neural decoding during encoding period (N =18).**

(A) Experiment 2 employed the same design as Experiment 1 (Figure 1A), except explicitly requiring participant to hold the ordinal position in WM. Specifically, during the recalling period, a target pure tone was presented, and participants needed to compare it to the cued memory tone, i.e., higher or lower in frequency (same as Exp 1), and then reported the ordinal position of the cued memory tone ($1^{st}$, $2^{nd}$, $3^{rd}$) by pressing corresponding keys (‘A’, ‘S’ and ‘D’). (B) Neural representations of content (i.e., frequency) and structure (i.e., ordinal position) during encoding period. Grand average (N = 18, mean ± SEM) beta values of the regression between neural representational dissimilarity and physical dissimilarity of memory tones (i.e., decoding performance), for frequency (left) and ordinal position (right), as a function of time. Gray bar on x-axis indicates memory tone presentation. (Shaded area represents ± 1 SEM across participants. ***: $p < 0.001$; **: $p < 0.05$; solid line: corrected using cluster-based permutation test, cluster-forming threshold $p < 0.05$).
